# T cells Mediate Progression of Load-Induced Osteoarthritis

**DOI:** 10.1101/2020.05.20.106435

**Authors:** Tibra A. Wheeler, Adrien Y. Antoinette, Matthew J. Kim, Marjolein C. H. van der Meulen, Ankur Singh

## Abstract

Osteoarthritis (OA) is a degenerative disease that manifests as joint damage and synovial inflammation. To date, most studies have focused on the decrease in cartilage stiffness, chondrocyte viability, and changes in matrix-degrading enzymes. With the exception of a few inflammatory cytokines and macrophages, the immune response in OA is poorly characterized, and the crosstalk of joint damage with T and B cells in local lymph nodes is unknown. Here, using an *in vivo* mouse model of mechanical loading of mouse tibia, we demonstrate that CD8+ T cells and subsets of CD4+ T cells, and not B cells, increase in the local lymph nodes and contribute to the progression of load-induced OA pathology. We demonstrate that T cell response is sex- and age-dependent. Mechanical loading of T cell knock-out mice that lack αβ T cell receptor carrying cells resulted in attenuation of both cartilage degradation and osteophyte formation in loaded joints, with a concomitant increase in γδ+ T cells. Restricting the migration of T cells in lymphoid tissues through the systemic treatment using Sphingosine-1-phosphate (S1P) inhibitor, decreased localization of T cells in synovium, and attenuated cartilage degradation. Our results lay the foundation of the role T cells play in the joint damage of load-induced OA and allude to the use of S1P inhibitors and T cell immunotherapies for slowing the progression of OA pathology.

## Introduction

Osteoarthritis (OA) is a progressive, degenerative, and disabling disease of articulating joints that affect approximately 27 million people in the US (*1–3*). Aging, obesity, genetics, occupational activities, and prior joint injury are all risk factors for OA, making it the leading cause of disability (*2, 3*). Repeated compressive cyclic loading of joints can decrease cartilage stiffness and chondrocyte viability with an increase in matrix metalloproteinases (MMPs) and aggrecanase expression (*4, 5*). Once the damage is evident radiographically, the progression is rapid and cannot be reversed. Therefore, there is a need to understand the mechanisms that participate in OA initiation and progression and develop new therapies.

Classically OA has not been viewed as an immune-based disease. Although the synovial membrane of OA patients has marked T cell infiltration, specifically CD4+ and CD8+ T cells (*6*), several standing questions remain in the field on the fate of T cells in local lymph nodes in load-induced OA and the impact of these T cells on the progression of OA pathology. We hypothesized that mechanical load-induced damage of joints initiates a bi-directional interaction between the lymph node and joint, and the immune response in the local lymph node will regulate OA progression. Here, using a combination of wild type and transgenic mice of multiple ages, we demonstrate unexpected increase in specific T cell populations in the regional lymph nodes in a mouse model of load-induced OA with single and multiple bouts of mechanical loading. Using a combination of T cell receptor knockout mice and WT mice treated with an inhibitor of T cell migration, we demonstrate that the change in T cells in lymph nodes and their migration was associated with OA pathogenesis in the daily cyclic tibial compression mouse model, as determined by cartilage damage and osteophyte formation. These findings correlated with the increase in T cells, and not B cells, in both mice and humans. These results highlight that a local lymph node is also a key modulator of the immune response in mechanical loading-induced OA, in contrast to the prior focus of knee joint only. Understanding the unique T cell immune response will enable a rational immunotherapeutic approach to be developed that can be designed to target specific lymphocyte populations in OA, for which no targeted therapy exists.

## Results and Discussion

### T cells in ipsilateral lymph nodes increased with cyclic tibial compression in female and male mice

We hypothesized that mechanical load-induced damage of joints initiates a bi-directional interaction between the lymph node and joint, localized to the loaded limb. To test our hypothesis, we induced OA pathology in wild type (WT, C57Bl/6) mice by applying cyclic compression (9.0N peak load, 1200 cycles, 4Hz frequency, 5 days/week) to the left tibiae of young (10-week-old) and adult (26-week-old) female mice. This loading regimen induces cartilage matrix damage, including proteoglycan loss, fibrillation, and erosion as well as osteophyte formation (*7–10*). In all studies, the contralateral limbs served as the control. We applied daily loading for one week and evaluated the lymphocyte counts using flow cytometry (**Figure 1A**). The size of the ipsilateral inguinal lymph nodes in the loaded limbs markedly increased compared to those in the contralateral limb of the same mouse (**Figure 1B**). The cyclic tibial compression increased the total number of CD3+, CD3+CD4+, and CD3+CD8+ T cells increased significantly in the inguinal lymph node of loaded limbs in 10 week (young) and 26 week (adult) old female mice (**Figure 1C**; **Supplementary Figure S1A**). To examine whether the T cell response was sex- or loading duration-dependent, we loaded adult WT male mice for 1 or 2 weeks. Unlike female mice, after one week of cyclic tibial compression, the total number of T cells did not change significantly in the ipsilateral inguinal lymph nodes compared to the contralateral limb. However, two weeks of loading significantly increased the total number of CD3+, CD3+CD4+, and CD3+CD8+ T cell subsets in the inguinal lymph node of the loaded limb (**Figure 1D**; **Supplementary Figure S1C**). This result is consistent with greater cartilage damage in the tibial plateau after two weeks than one week of cyclic tibial loading in adult male mice (*10*). These findings further allude to the time dependency of the T cell response with the progression of OA.

**Fig 1.**
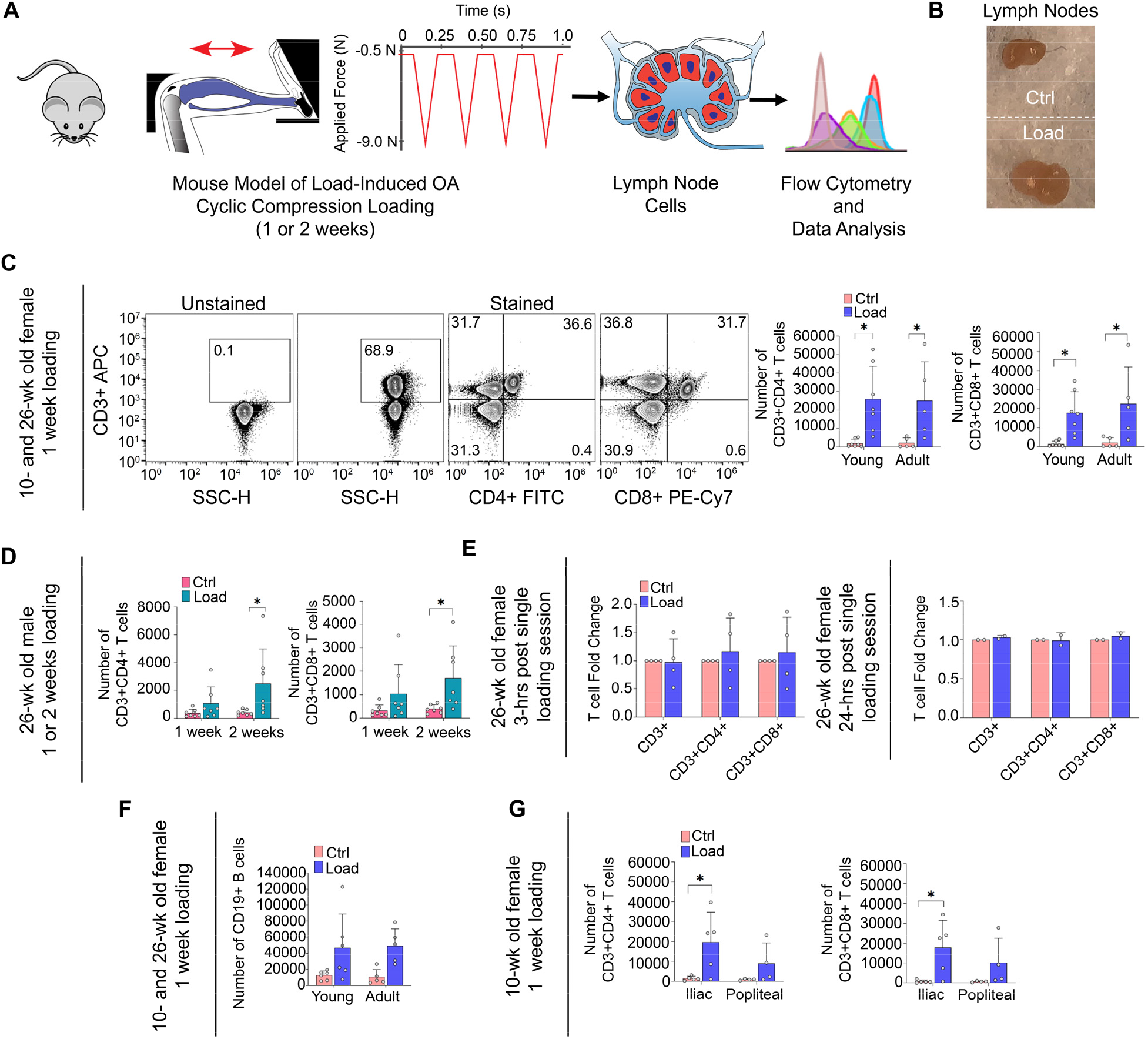
Abundance of T cells increased in inguinal lymph nodes after 1 or 2 weeks of cyclic tibial compression in male and female mice. **A)** Mouse cyclic tibial compression model. Schematic of tibia positioned in loading device with 9N peak load magnitude. Schematic of experimental design. Mice underwent 1 or 2 weeks of cyclic tibial compression and the inguinal lymph node was assessed through flow cytometry. **B)** Images of inguinal lymph nodes corresponding to the contralateral (left) and loaded (right) limbs. **C)** Gating strategy for CD3+, CD3+CD4+, and CD3+CD8+ T cells showing the percentage of the T cell markers after 1 week of loading in 10-and 26-week old female C57Bl/6 mice. Comparison of the number of CD3+CD4+, and CD3+CD8+ T cells in the inguinal lymph node of loaded and contralateral limbs for female mice. *p<.05 **D)** Comparison of the number of CD3+CD4+, and CD3+CD8+ T cells in the inguinal lymph node of loaded and contralateral limbs for male mice. *p<.05 **E)** Fold change of loaded relative to contralateral in CD3+, CD3+CD4+, and CD3+CD8+ T cell markers 3- and 24-hours post-loading in 26-week old C57Bl/6 female mice. **F)** Comparison of the number of CD19+ B cells in the inguinal lymph nodes of loaded and contralateral limbs for 10- and 26-week old C57Bl/6 female mice. *p<.05 **G)** Comparison of the number of CD3+CD4+ and CD3+CD8+ T cells in the iliac and popliteal lymph nodes of loaded and contralateral limbs after 1 week of loading in 10-week old female C57Bl/6 mice. *p<.05

To understand the timing of the immune response, we examined 3- and 24-hours after the initiation of loading in adult female mice. At these early time points, the number of T cells in the inguinal lymph nodes were not different between loaded and contralateral limbs (**Figure 1E**). These findings suggest that the increase in T cells is a temporal process developing after the initiation of load-induced damage. Interestingly, the increase in lymphocytes was only significant in T cells and CD19+ B cell numbers did not increase in the ipsilateral inguinal lymph nodes as compared to contralateral side (**Figure 1F**). Because the knee drains to multiple lymph nodes (*11*), we further compared the popliteal and iliac lymph nodes. Following *in vivo* loading, all ipsilateral lymph nodes showed an increase in T cell numbers, with iliac and inguinal being similar in response and popliteal to a lesser extent (**Figure 1G**; **Supplementary Figure S1B**). We focused on the inguinal lymph node because the T cell response was slightly higher than the iliac.

### Macrophage population increased with cyclic tibial compression in sex- and age-dependent manner

Macrophages are associated with inflammation in OA (*12*). Using 10- and 26-week-old C57Bl/6 female mice, we examined whether the macrophage cell population changed with cyclic tibial compression of the left tibia (9.0N peak, 1200 cycles, 4Hz frequency, 5 days/week). Using flow cytometry, the number of F4/80+ macrophages increased in the inguinal lymph node of the loaded limb of only adult mice after a week of loading compared to young female mice (**Figure 2A**). Within the increased macrophage population we further investigated whether loading induced differences in M1 and M2 extreme polarized macrophages phenotypes (*13*). Previously, a mixture of both M1- and M2-related genes was expressed by synovial macrophages across OA patients (*12*). Interestingly, both the number of M1 (CD197+) and M2 (Egr2+) macrophage subsets increased significantly with tibial loading in the inguinal lymph node of adult female mice (**Figure 2A**). Our results suggest differential activities of macrophages within the extremes of the M1/M2 spectrum. Whereas M1 macrophages may contribute to the inflammation with repetitive loading, the increased M2 macrophages may indicate inflammatory resolution. In contrast, we observed no significant changes in the total number of macrophages in the inguinal lymph nodes at 3- and 24-hours after initiation of loading (**Supplementary Figure S2**). This time-dependent response of the macrophage points to a chronic inflammatory response. The macrophage response was sex-dependent for the ages and loading durations studied. Unlike their female counterparts, inguinal lymph nodes of adult male mice had no significant differences in macrophage numbers after 1 or 2 weeks of repetitive loading compared to the contralateral control (**Figure 2B**). The gender dependency could be attributed to higher prevalence of macrophages in women than men with OA (*14*). Furthermore, within female mice, this response was age dependent.

**Fig 2.**
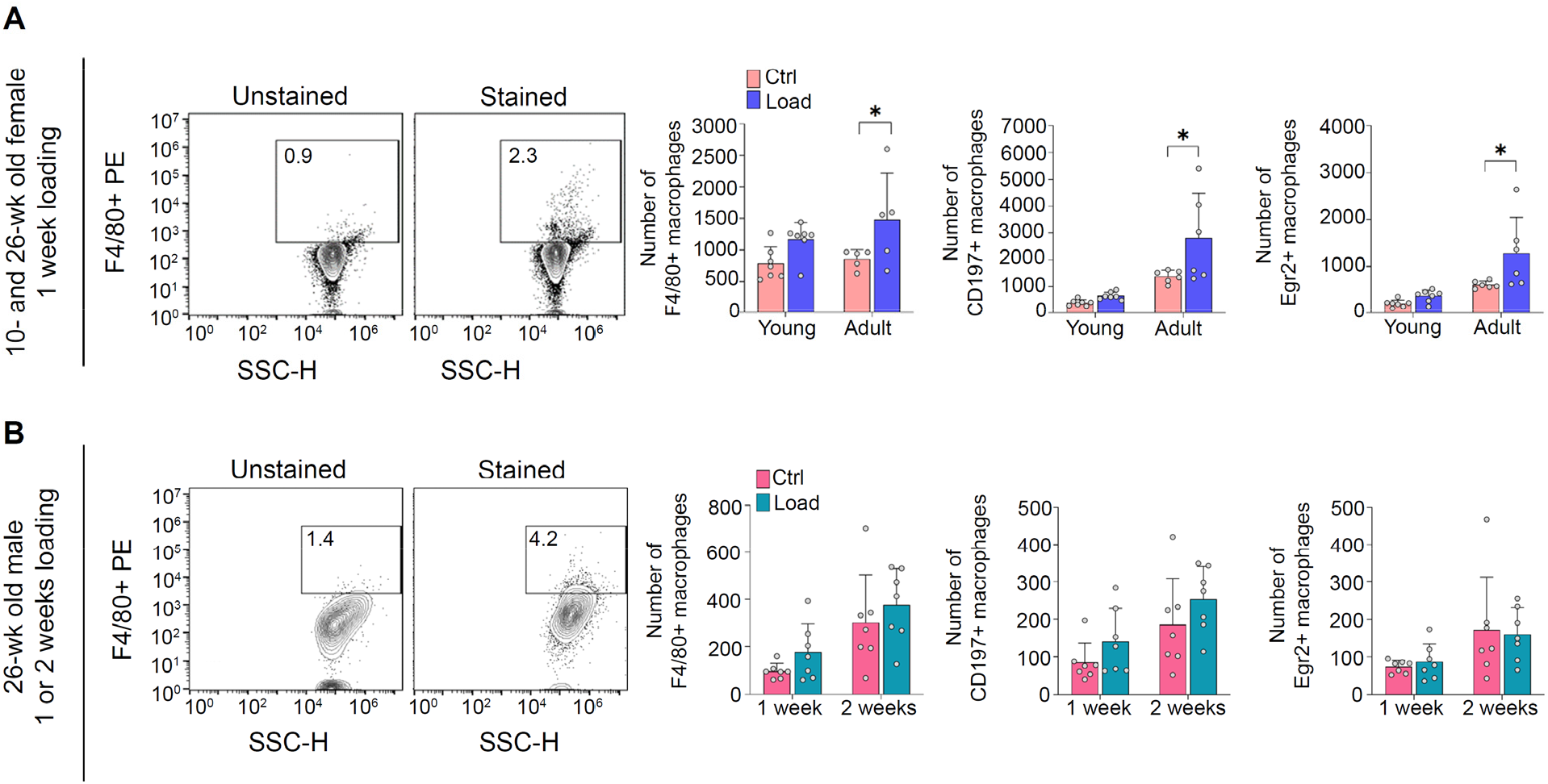
Abundance of macrophages increased in inguinal lymph nodes after 1 or 2 weeks of cyclic tibial compression in adult female mice. **A)** Gating strategy for F4/80+ macrophages showing the percentage of the macrophage marker after 1 week of loading in 10- and 26-week old female C57Bl/6 mice. Comparison of the number of F4/80+ (pan marker), CD197+ (M1), and Egr2+ (M2) macrophages in the inguinal lymph node of loaded and contralateral limbs for female mice. *p<.05. **B)** Gating strategy for F4/80+ macrophages showing the percentage of the macrophage marker after 1 or 2 weeks of loading 26-week old male C57Bl/6 mice. Comparison of the number of F4/80+ (pan marker), CD197+ (M1), and Egr2+ (M2) macrophages in the inguinal lymph node of loaded and contralateral limbs for male mice. *p<.05.

### Number of T cells and macrophages increased with cyclic tibial compression in mice lacking Deletion of estrogen receptor-alpha in osteoblasts

Women are more prone to knee OA than men, particularly post-menopausal women who are estrogen-deficient (*15*). To understand whether the increases in both T cells and macrophages in female mice were estrogen-dependent, we induced load-based OA pathology in 16-week-old female transgenic ERαKO mice lacking estrogen receptor-α in mature osteoblasts and littermate WT controls (*16, 17*). We chose 16-week-old mice because load-induced damage and OA development in 10- and 26-wk age mice were similar (*10*), and the T cell response in female mice was not age-dependent (Figure 1). After 2 weeks of *in vivo* tibial loading, more severe joint damage was present in the pOC-ERaKO mice compared to littermate controls (*16*). The numbers of T cells (CD3+CD4+ and CD4+CD8+) and F4/80+ macrophages increased in the inguinal lymph nodes of the loaded limbs, in the absence of ER-α in osteoblasts and their littermate controls (**Figure 3**).

**Fig 3.**
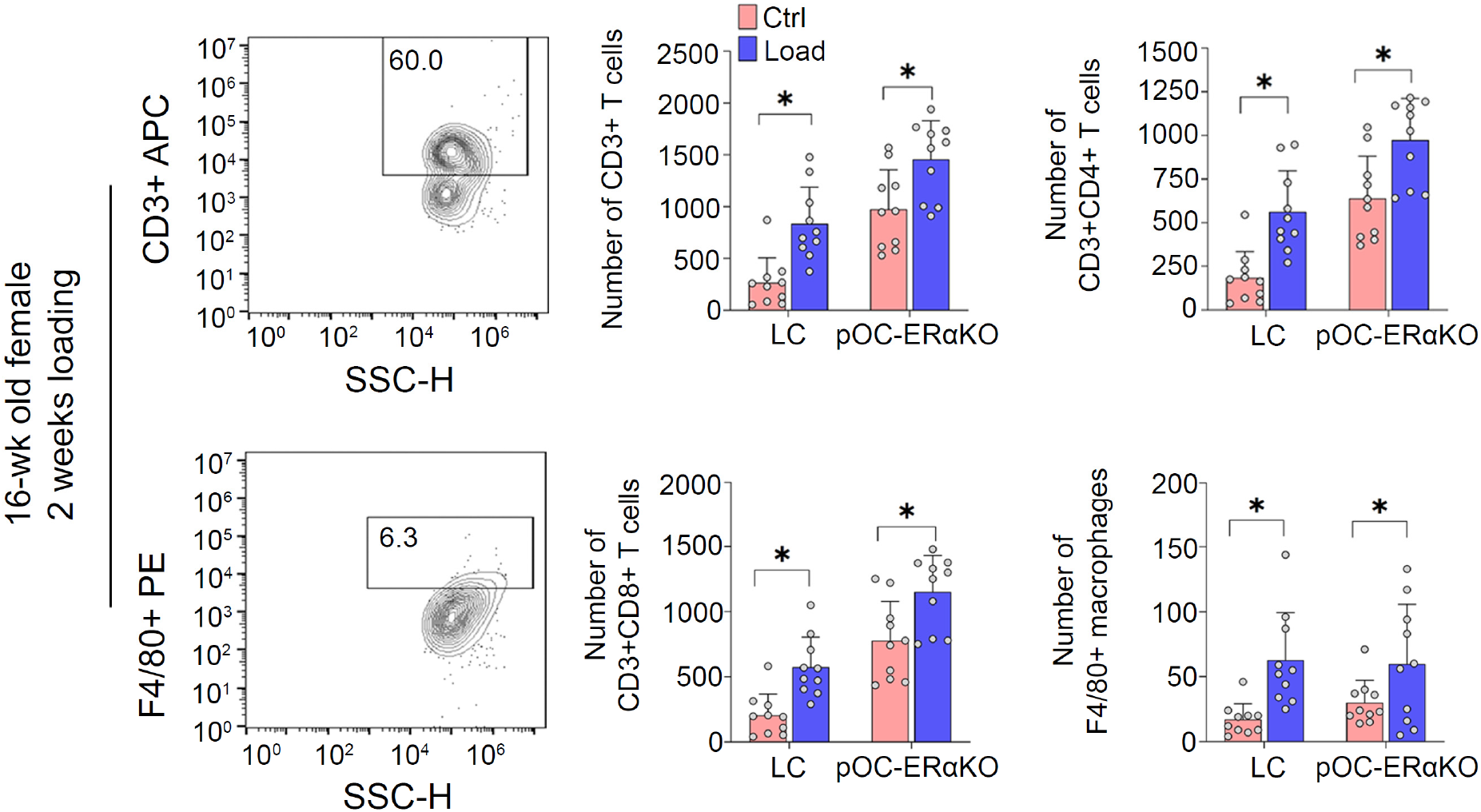
Number of T cells and macrophages increased in the inguinal lymph node after 2 weeks of cyclic tibial compression in skeletally mature pOC-ERαKO mice and littermate controls. Gating strategy for CD3+, CD3+CD4+, and CD3+CD8+ T cells and F4/80+ macrophages showing the percentage of these markers after 2 weeks of loading in 16-week old female LC and pOC-ErαKO mice. Comparison of the number of CD3+, CD3+CD4+, and CD3+CD8+ T cells and F4/80+ macrophages in the inguinal lymph node of loaded and contralateral limbs for female LC and pOC-ErαKO mice. *p<.05.

These findings allude to the fact that an increase in immune cell numbers in the inguinal lymph nodes and the differences between female and male mice were not simply attributable to estrogen signaling and additional factors, including anatomical differences and microbiome, remain to be investigated.

### Fingolimod treatments did not change the number of T cells with cyclic tibial compression

With the increase in T cells in the local lymph node following damaging mechanical loading, we hypothesized that T cells play a role in the progression of load-induced OA. We first sought to understand whether the migration of T cells out of the inguinal lymph node affected the progression of load-induced OA. Previous studies have shown that the immunomodulatory agent Fingolimod (FTY720) affects the ability of lymphocytes, specifically T cells, to respond to sphingosine‐1‐phosphate (S1P) through S1P receptors, ultimately affecting their ability to traffic through major lymphoid organs (*18, 19*). We applied loads to the knee joints of 16-week-old C57Bl/6 female mice for two weeks and simultaneously administered daily intraperitoneal (i.p.) injections of Fingolimod (1mg/kg) starting the first injection 24 hours before initiating loading. A single administration of fingolimod reduces peripheral lymphocyte counts within a few hours (*20*); however, the reduction is transient, and normal lymphocyte values return one day later. To determine whether specific T cell subsets increased disproportionately over others, we analyzed the inguinal lymph node using multi-color flow cytometry (**Figure 4A**). Fingolimod treatment increased the lymph node size in both contralateral and loaded limbs (**Figure 4B**), likely because of the restricted egress of T cells. The tissue-level response of the joint was analyzed by histology, microCT, and immunohistochemistry. Based on quantitative flow cytometry analysis two weeks of cyclic tibial compression upregulated specific T cell subsets in the loaded limbs of saline-treated mice compared to contralateral limbs. Specifically, the total numbers of CD3+, CD3+CD4+, CD3+CD4+forkhead box p3 (FoxP3)+ regulatory T (Tregs) cells, and γδ T cell receptor (TCR)+ T cell subsets increased significantly in the inguinal lymph node of loaded limbs; the number of CD3+CD8+ T cells also trended toward an increase. The increase in Tregs was intriguing because these cells can be anti-inflammatory. In recent studies, the profile of Tregs in OA patients was similar to that of rheumatoid arthritis patients with decreased expression and function of these cells (*21, 22*). However, the frequency of T regulatory cells in the blood of OA patients was elevated (*21*). We postulate that *in vivo* loading may increase the number of Treg cells, but further analysis of their function is needed. Interestingly, the number of CD3+CD4+GATA3+ (T_H_2) and CD3+CD4+Tbet+Rorγt+ T cell (T_H_17) subsets did not change in the inguinal lymph node with *in vivo* loading (**Figure 4C**; **Supplementary Figure S3**). T_H_2 and T_H_17 cells play a limited role in the pathogenesis of OA and show little alteration in the peripheral blood, synovial fluid, and synovial membranes of OA patients (*23*). Fingolimod treatment did not affect any of the T cell subsets in the inguinal lymph node of the loaded limb compared to contralateral controls (**Figure 4C**; **Supplementary Figure S3**). No increase in lymph node size is not surprising because at any time, only 2% of the total lymphocyte population circulates in the peripheral blood (*24*), explaining the lack of cell accumulation or abnormal lymph node enlargement with Fingolimod treatment.

**Fig 4.**
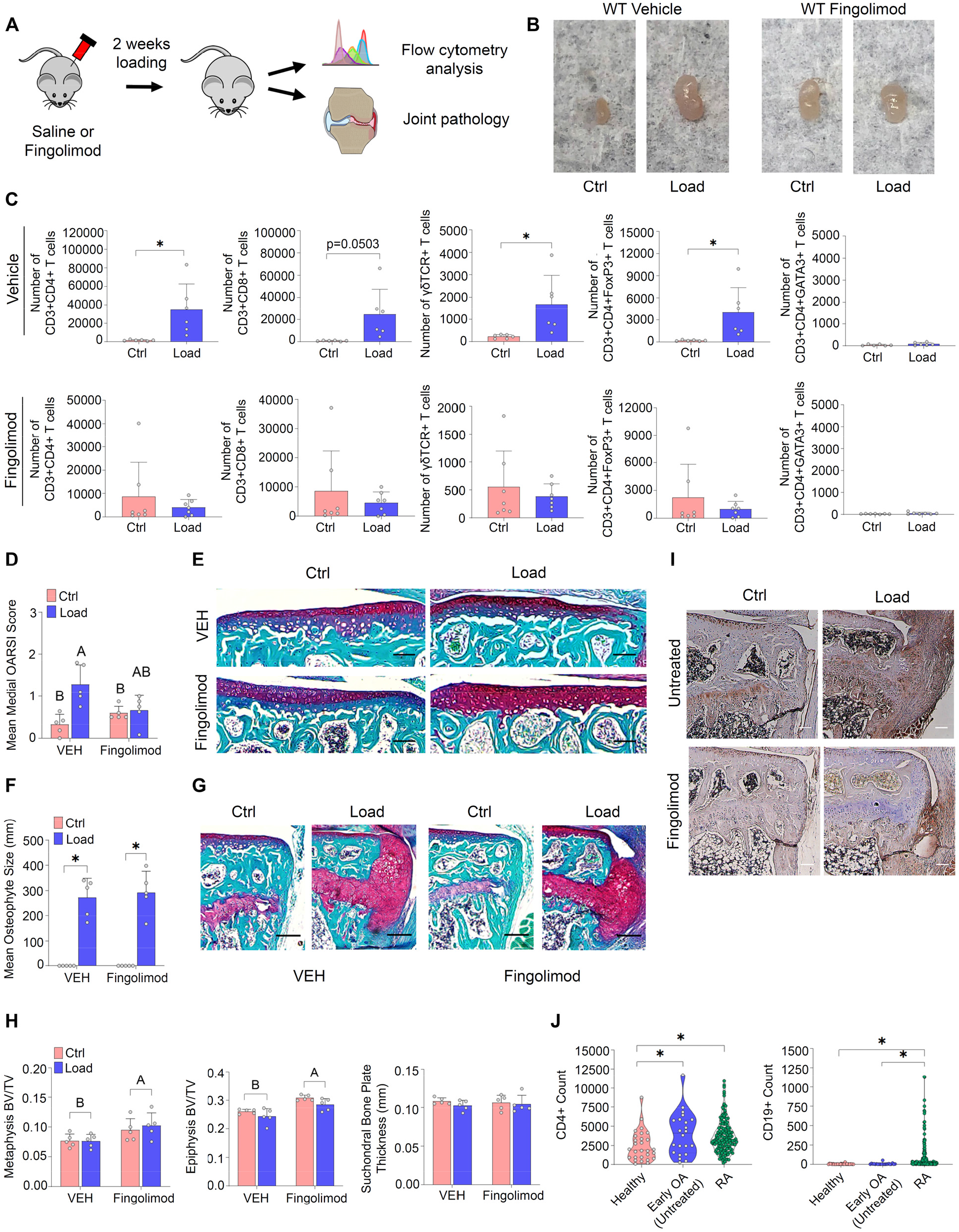
Fingolimod treatment attenuated load-induced cartilage degradation but did not change the number of T cells in the inguinal lymph node after 2 weeks of cyclic tibial compression. **A)** Schematic of experimental design. 16-week old female C57Bl/6 mice underwent 2 weeks of cyclic tibial compression with daily saline or Fingolimod i.p. injections. The inguinal lymph node was assessed through flow cytometry and joints were assessed through histology and microCT. **B)** Images of inguinal lymph nodes corresponding to the contralateral and loaded limbs for both the saline and Fingolimod treated groups. **C)** Comparison of the number of CD3+CD4+, CD3+CD8+, γδTCR, CD3+CD4+FoxP3+, and CD3+CD4+GATA3+ T cells in the inguinal lymph node of loaded and contralateral limbs of 16-week old female saline and fingolimod treated C57Bl/6 mice after 2 weeks of loading. *p<.05 **D)** Comparison mean OARSI scores in the medial tibial plateau after 2 weeks of loading and fingolimod treatment *p<.05 **E)** Safranin-O/Fast Green images of the contralateral and loaded limbs indicating cartilage erosion; scale bar = 100μm **F)** Comparison osteophyte size in the medial tibial plateau after 2 weeks of loading and fingolimod treatment *p<.05 **G)** Safranin-O/Fast Green images of the contralateral and loaded limbs indicating osteophyte formation; scale bar = 100μm **H)** microCT analysis comparison of subchondral bone plate thickness and epiphysis and metaphysis BV fraction after 2 weeks of loading and fingolimod treatment. A>B>C **I)** IHC images of the contralateral and loaded limbs indicating CD3+ T cell staining; scale bar = 100μm **J)** Comparison RNA sequencing performed on samples isolated from joint synovial biopsies (GSE89408) *p<.05

Similar to the T cells, the macrophage populations in the saline and Fingolimod treated mice increased with repetitive loading. The total numbers of F4/80+, F4/80+CD197+, and F4/80+Egr2+ macrophages increased significantly in lymph nodes of loaded limbs of saline-treated mice. Furthermore, loading increased CD169+ macrophages (**Supplementary Figure S4**), a specialized subtype of tissue-resident macrophages (*25, 26*). Fingolimod treatment did not change the number of macrophage subsets in the inguinal lymph node of the loaded limb compared to contralateral controls (**Supplementary Figure S4**), which may reflect that macrophages and monocytes express S1P receptors (*27*).

To determine the effects of restricting T cell migration on load-induced OA, we analyzed knee joints after two weeks of tibial loading and treatment. Key hallmarks of our load-induced OA model are cartilage degradation, osteophyte formation, and changes in bone morphology including epiphyseal and metaphyseal bone adaptation. In general, loading induced cartilage erosion extending to the superficial layer of the tibial cartilage. The loaded limb of saline-treated mice had significantly increased OARSI histology scores indicating cartilage damage compared to contralateral controls. However, with Fingolimod treatment, the cartilage in the loaded limb was not different compared to contralateral controls (**Figure 4D, E**), suggesting a role of T cell migration in cartilage degradation. In both saline- and Fingolimod-treated groups, osteophytes formed on the medial tibial plateau with two weeks of loading; osteophytes were not present in contralateral control joints (**Figure 4F, G**; **Supplementary Figure S5**).

Loading did not alter subchondral bone plate thickness or tissue mineral density (TMD) in either treatment group. However, TMD was increased in Fingolimod-treated mice compared to saline-treated mice in both loaded and contralateral limbs. Subchondral bone plate thickness was not different between treatment groups. Similarly, the metaphyseal and epiphyseal cancellous BV/TV were higher in Fingolimod-treated mice compared to saline-treated mice, but not affected by loading in either treatment group (**Figure 4H**; **Supplementary Figure S5**).

Based on the absence of OA-like pathology with combined loading and Fingolimod treatment, we hypothesized that migratory, not tissue-resident, T cells play a role in load-induced OA. Therefore, we performed immunohistochemistry for CD3+ T cells in the joints of mice after loading treated with Fingolimod or saline. In the saline-treated mice, loaded joints demonstrated a high number of T cells in the osteophyte and synovium regions. In contrast, tissues from Fingolimod-treated mice had markedly less CD3+ T cell staining in the joint (**Figure 4I**). To further validate the presence of T cells in the synovial joints of human OA patients, we examined RNA sequencing data (GSE89408) from synovial biopsies of patients with normal, early OA, and rheumatoid arthritis reported by Guo et al (*28*). CD4 gene expression counts increased between the normal tissues and early OA samples. The increase in T cell genes with OA was similar to that of tissue from rheumatoid arthritis patients. As expected, the CD19 (B cell) gene expression counts were not different between the normal tissues and the early OA tissues, and significantly lower than CD19 counts in rheumatoid arthritis patients (**Figure 4J**).

### Absence of αβTCR+ T cells, but not γδTCR+ T cells, attenuated load-induced cartilage degradation and osteophyte size

Our results from T cell migration restriction suggested that migratory T cell subsets may be playing a specific role in load-induced OA. In particular, γδTCR+ T cells emerged as a critical non-CD4+ T cell subset. To determine whether γδTCR+ T cells played a role in OA, we induced load-induced OA in 16-week-old TCRα-lacking transgenic C57Bl/6 mice (*29*). The TCRαKO homozygous mice are devoid of CD4+CD8− and CD4-CD8+ T cells. These mice, however, have functional γδ T cells, a subset of T cells with distinct TCR γ and δ chains on their surface that account for 0.5–5% of all T cells in a mouse. Comparisons were made to WT (C57Bl/6), the background strain for TCRα −/− mice. After two weeks of cyclic tibial compression, the inguinal lymph node and knee joints were analyzed (**Supplementary Figure S6A**). While the total numbers of CD3+ T cells, and CD3+CD4+ and CD3+CD8+ T cell subsets increased significantly in the inguinal lymph nodes of the loaded limbs in WT mice, we did not observe an increase in the TCRα −/− mice. However, the total number of γδTCR+ T cell subset significantly increased with loading in the TCRα −/− mice (**Figure 5A**, **Supplementary Figure S6B**). Furthermore, two weeks of loading increased the total number of F4/80+ macrophages in both WT and TCRα −/− mice. Interestingly, the increase in CD197+ (M1) and Egr2+ (M2) macrophage subsets was only significant in TCRα −/− mice, as these numbers remained indifferent between the contralateral and loaded limb inguinal lymph nodes of WT mice (**Supplementary Figure S6C**). However, whether this increase plays a role in OA is unclear.

**Fig 5.**
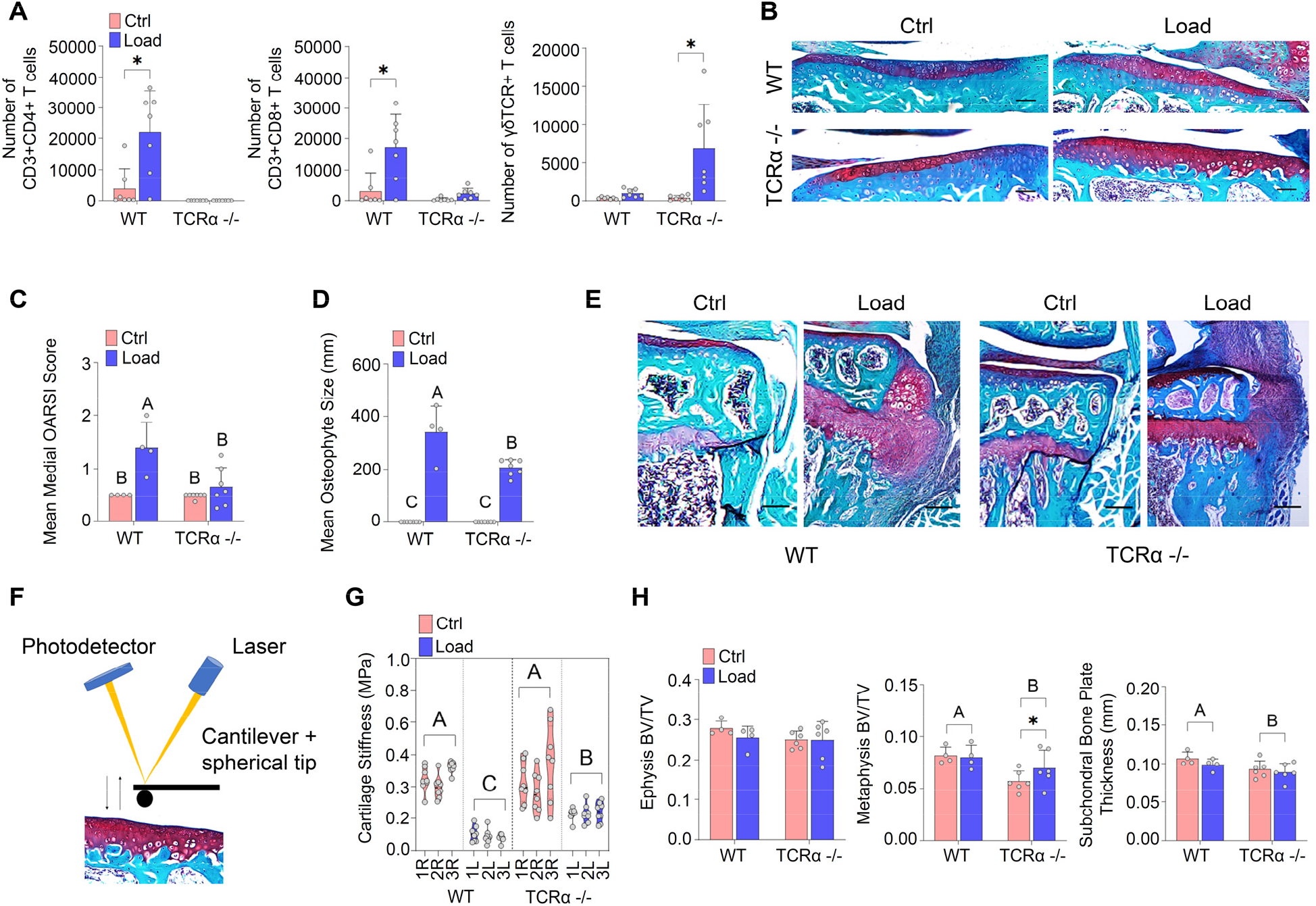
The absence of αβ T cells attenuated load-induced cartilage degradation and osteophyte size. **A)** Comparison of the number of CD3+CD4+, CD3+CD8+ and γδTCR T cells in the inguinal lymph node of loaded and contralateral limbs of 16-week old female C57Bl/6 and TCRα −/− mice after 2 weeks of loading. **B)** Safranin-O/Fast Green images of the contralateral and loaded limbs indicating cartilage erosion; scale bar = 100μm **C)** Comparison mean OARSI scores in the medial tibial plateau in 16-week old female C57Bl/6 and TCRα −/− mice after 2 weeks of loading. A>B **D)** Comparison osteophyte size in the medial tibial plateau in 16-week old female C57Bl/6 and TCRα −/− mice after 2 weeks of loading. A>B>C **E)** Safranin-O/Fast Green images of the contralateral and loaded limbs indicating osteophyte formation; scale bar = 100μm **F)** Schematic of AFM-nanoindentation on the tibial cartilage. **G)** Comparison of cartilage indentation stiffness in 16-week old female C57Bl/6 and TCRα −/− mice after 2 weeks of loading. A>B>C **H)** microCT analysis comparison of subchondral bone plate thickness and epiphysis and metaphysis BV fraction in 16-week old female C57Bl/6 and TCRα −/− mice after 2 weeks of loading. A>B

To determine the effects of the absence of αβTCR+ T cells, an increase in γδTCR+ T cells, and an unexpected increase in macrophages (M2 > M1) on load-induced OA, we analyzed knee joints after two weeks of mechanical loading. When we compared the loaded limb to contralateral controls of WT mice, the loaded limbs had increased cartilage damage scores compared to control limbs. In contrast, cartilage damage in the loaded limb was not different when compared to contralateral control limbs of TCRα −/− mice (**Figure 5B, C**), suggesting a role of T cells in cartilage degradation. In all groups, osteophytes formed on the medial tibial plateau with two weeks of loading. Osteophyte maturity was similar in WT and TCRα −/− mice; however, the osteophytes in the TCRα −/− mice were significantly smaller than those seen in WT mice, further suggesting a role of T cells in the presence or growth of osteophytes (**Figure 5D, E**; **Supplementary Figure S7A**).

We next sought to determine whether the lack of cartilage damage in the TCRα −/− mice was attributable to altered cartilage mechanical properties as compared to WT mice. We measured the nanomechanical properties of cartilage relative to the development of OA using atomic force microscopy (AFM)-based nanoindentation on tibial condyle cartilage surfaces, as reported elsewhere (*30*) (**Figure 5F**). Two weeks of loading reduced cartilage stiffness in the loaded limbs compared to contralateral controls of WT mice and TCRα −/− mice; however, the overall stiffness of the loaded limb cartilage was reduced less in TCRα −/− mice than WT mice (**Figure 5G**).

Loading did not affect the subchondral bone plate thickness and TMD or epiphyseal cancellous BV/TV; however, WT mice had overall thicker subchondral bone plates and metaphyseal cancellous BV/TV compared to the TCRα −/− when both contralateral and loaded limbs were pooled (**Figure 5H**; **Supplementary Figure S7B**). TMD and epiphyseal cancellous BV/TV were not different between genotypes. Unlike the subchondral plate, loading increased metaphyseal cancellous BV/TV, but only in the TCRα −/− mice. Even with thinner subchondral bone plates and lower metaphyseal BV/TV, the TCRα −/− mice demonstrated improved cartilage health after loading indicating that the bone properties were not a contributing factor.

## Conclusion

Classically OA has not been viewed as an immune-based disease and no previous studies have elucidated the role of load-induced OA on the lymph nodes and the impact of the local presence of T cells on the progression of joint tissue damage in load-induced OA. Given the emerging need for new OA therapies, understanding the immune response will enable the development of a rational immunotherapeutic approach to target specific immune cells. We demonstrated that the local lymph nodes are key modulators of the immune response in load-induced OA. Collectively, based on our findings that mice with normal levels of T cells in lymph nodes had more severe OA pathology, increased localization of T cells in synovial joints, and reduced cartilage stiffness compared to the absence of T cells, indicating T cells play a crucial role in cartilage degradation and progression of OA. The T cell depletion and lack of trafficking studies used skeletally mature female mice. However, females and younger mice tend to have less severe OA so further investigation is needed to confirm the gender and age-dependency of the T cell response. We focused on repetitive bouts of loading, which mimic the development of OA over time; however, joint injuries lead to post-traumatic OA, which may have a different outcome. Further studies are needed to elucidate the role of T cells after a single bout of loading beyond the 3-hour time point reported here. We anticipate that these studies lay the foundation for the role T cells play in joint damage associated with load-induced OA and further allude to the use of S1P inhibitors and immunomodulators as potential therapeutics for the treatment of OA.

## Materials and Methods

### Mice

C57Bl/6 male and female mice (10-, 16-, and 26-week old) were purchased from Jackson Laboratories (n=5-7 per group). For estrogen receptor-α conditional knockout experiments, female mice were generated using the OC-Cre to remove ERα at the stage of mature osteoblasts and osteocytes (pOC-ERαKO). The pOC-ERαKO and littermate control (LC) mice were bred and validated as previously described (*17*). For T cell knockout experiments, we used TCRα-lacking transgenic C57Bl/6 mice (purchased from Jackson Laboratories and gifted by Brian Rudd Lab at Cornell). To inhibit T cell migration, Fingolimod (FTY720, Cayman Chemical Co) was administered to C57Bl/6 mice. Administration of Fingolimod or saline vehicle was performed by daily intraperitoneal injections (1mg/kg BW i.p.) with the first dose administered 24 hours before loading (*31, 32*). All experimental techniques were approved by the Cornell Institutional Animal Care and Use Committee.

### In vivo mechanical loading

To induce OA, we applied cyclic compression to the left tibia of mice at a 9.0N-peak magnitude for 1200 cycles at a 4Hz frequency; contralateral limbs served as controls(*10*). Female WT mice were euthanized 3 hours after a single session of loading or after 1 or 2 weeks of daily loading (5 days/week); male WT mice were euthanized after 1 or 2 weeks of daily loading (5 days/week). TCRα-lacking and pOC-ERαKO mice were euthanized after 2 weeks of daily loading (5 days/week).

### Flow cytometry

Inguinal LNs were removed and dissociated into single-cell suspensions. The suspensions were divided into equal volumes for analysis. Cells were stained for cell surface and intracellular markers listed in Supplementary Table 1. Extracted cells were stained by incubation in FACS buffer containing antibodies with 1:200 dilution for 1 hour in the dark at 4°C. The entire volume was analyzed for the total number of cells (BD Accuri C6; BD Biosciences Symphony Analyzer; FlowJo).

### Bone morphological changes

Knee joints were harvested and fixed in 4% paraformaldehyde and scanned in 70% ethanol at 15μm voxel resolution (μCT35, Scanco, Bruttisellen, Switzerland; 55 kVp, 145mA, 600 ms integration time), as reported earlier(*9, 17*). Cancellous and cortical bone in the epiphysis and metaphysis of the proximal tibia were analyzed to assess bone morphology using microcomputed tomography (microCT). The subchondral bone plate was analyzed for thickness and tissue mineral density (TMD). The cancellous bone in the epiphysis and metaphysis were analyzed for bone volume fraction (BV/TV).

### Cartilage degradation

After microCT scanning, tissues were decalcified in 10% ethylenediaminetetraacetic acid (EDTA) and processed for paraffin embedding for analyzing cartilage degradation(*10, 33*). Paraffin blocks were sectioned from posterior to anterior at 6-μm thickness using a rotary microtome (Leica RM2255, Wetzlar, Germany). Sections were stained using Safranin-O/Fast Green at 90-μm intervals to assess cartilage morphology in the medial tibial plateau. The modified OARSI scoring system was used to histologically assess cartilage damage. Mean scores were calculated for each limb (*34*).

### Osteophyte formation

To assess osteophyte formation, Safranin-O/Fast Green histological sections were examined by analyzing medial osteophytes in the medial tibial plateau from three representative sections in the joint (anterior, middle, and posterior). Osteophyte maturity(*35*) was evaluated based on the degree of calcification of ectopic bone. Osteophyte size was based on the medial-lateral width, defined as the distance between the medial end of the epiphysis and the end of the osteophyte; the mean width was calculated for each limb (*9*).

### Immunohistochemistry

T cell (CD3+) expression was assessed using IHC. Representative sections from the middle region of the joint were stained. Sections were deparaffinized, rehydrated, incubated in citrate retrieval buffer at 60°C for 60 min, and incubated with 0.3% H_2_O_2_ solution for 10 min. Immunostaining was performed using a rabbit specific HRP/DAB (ABC) IHC detection kit (ab64261, Abcam). Sections were blocked with protein block for 5 min and incubated overnight at 4°C with either rabbit polyclonal anti-CD3 antibody (1:50 dilution, ab5690, Abcam) or rabbit monoclonal negative control anti-IgG (1:50 dilution, ab172730, Abcam). Sections were treated with biotinylated goat anti-rabbit IgG (H+L) for 10 min, followed by treatment streptavidin peroxidase for 10 min. Finally, sections were incubated with DAB chromogen solution (1:50 dilution) for 5 min, counterstained with hematoxylin for 1 min, and mounted with a coverslip.

### Nanoindentation

Nanoindentation was used to assess cartilage mechanical properties. Knee joints were harvested and frozen in PBS-soaked gauze. Limbs were defrosted, and the femur, ligaments, tendons, and meniscus were dissected to expose the tibial cartilage. Tibial plateaus with exposed cartilage were then glued to the surface of a microscope slide (Loctite 409, Henkel Corp., Rocky Hill, CT). During indentation, tibial cartilage was maintained in PBS to prevent dehydration. Atomic force microscope (AFM)-based nanoindentation was performed on the medial and lateral tibial condyles with a spherical tip (Borosilicate particle, 5-μm radius, Au coating, Novascan Technologies) using an AFM (Asylum-MF3D-Bio-AFM-SPM), as previously described (*30*); indentation was repeated at 7-9 different locations per tibia.

### Statistical analysis

Statistical analysis was performed as detailed in the figure legends and text. All lymph node, bone morphology, and joint pathology analyses used a two-factor analysis of variance (ANOVA) with Sidak’s test, a paired two-tailed t test, or a linear mixed-effects model with fixed effects of loading and genotype or treatment with Tukey’s post hoc. Nanoindentation experiments used a two-factor ANOVA with Tukey’s post hoc test. Quantitative analyses as bar graphs with individual points are presented as means + SD. Quantitative analyses as violin plots with individual points, dotted lines represent upper and lower quartiles and median. Each datapoint represents a single mouse. In all studies, *p<0.05. Lack of significance is denoted by “ns.” On plots with letters, bars that share a letter are not significantly different from each other, A>B>C.

## Acknowledgments

We thank Dr. Lionel Ivashkiv and Hayat Ben Larbi at the Hospital for Special Surgery, Chiemezue Ijomanta, Simon Ortiz, Carolyn Chlebek, Amanda Rooney, Kristine Lai, Dr. Brian Rudd and his laboratory, Dr. Glenn Jackson, and the Center for Animal Resources and Education (CARE) staff at Cornell University. We acknowledge partial financial support from the 3M Non-Tenured Faculty Award (AS), National Institutes of Health (R21-AR064034, MvdM), Department of Defense (USAMRMC W81XWH-17-1-0540, MvdM), National Science Foundation (#1605935, MvdM), National Science Foundation Graduate Research Fellowship (DGE-1650441, TAW), and the Cornell Sloan and Colman Diversity Fellowships (TAW).

## Author Contributions

Concept and design: TAW, MvdM, AS

Acquisition: TAW, AYA, MJK, MvdM, AS

Analysis and interpretation of data: TAW, AYA, MJK, MvdM, AS

Drafting and critical revision of article: TAW, MvdM, AS

Final approval of article: TAW, AYA, MJK, MvdM, AS

## Notes

### Competing Interest Statement

The authors have declared no competing interest.

