## Supplementary Figures for "T cells Mediate Progression of Load-Induced Osteoarthritis"

Supplementary Figure S1

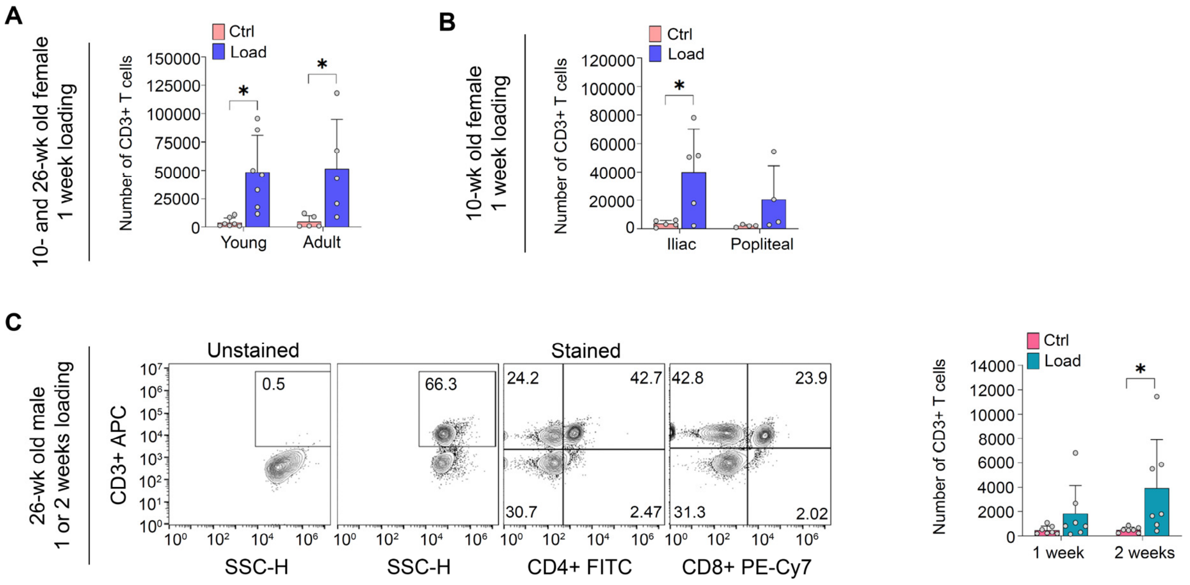


**Supplementary Figure S1, Related to Figure 1. Abundance of T cells and macrophages increased in inguinal lymph nodes after one or two weeks of cyclic tibial compression in female mice. A)** Comparison of the number of CD3+ T cells in the inguinal lymph nodes of loaded and contralateral limbs for 10- and 26-week old C57Bl/6 female mice. *p<.05 **B)** Comparison of the number of CD3+T cells in the iliac and popliteal lymph nodes of loaded and contralateral limbs after 1 week of loading in 10-week old female C57Bl/6 mice. *p<.05 **C)** Gating strategy for CD3+, CD3+CD4+, and CD3+CD8+ T cells showing the percentage of the T cell markers after one or two weeks of loading in 26-week old male C57Bl/6 mice. Comparison of the number of CD3+ T cells in the inguinal lymph node of loaded and contralateral limbs for male mice. *p<.05

Supplementary Figure S2

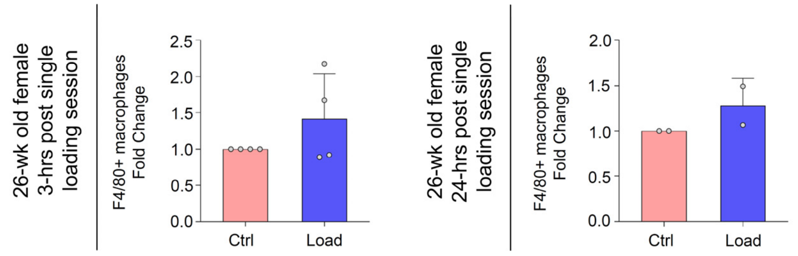


**Supplementary Figure S2, Related to Figure 1. Abundance of macrophages did not change in inguinal lymph nodes 3 or 24 hours after a single session of cyclic tibial compression in adult female mice.** Fold change in F4/80+ macrophages 3- and 24-hours post-loading in 26-week old C57Bl/6 female mice.

Supplementary Figure S3

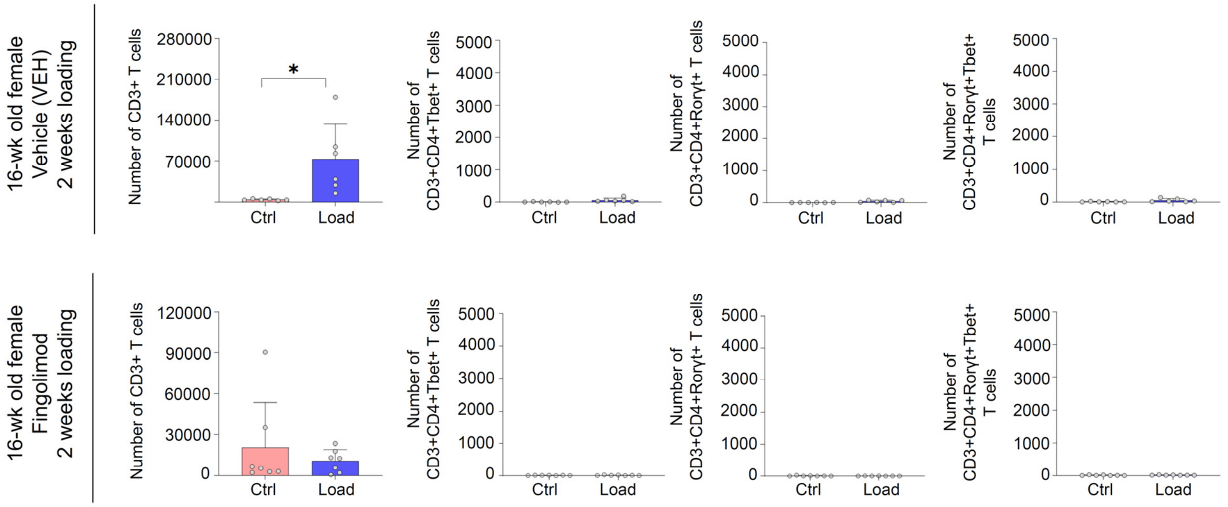


**Supplementary Figure S3, Related to Figure 4. Fingolimod treatment did not change the number of T cells in the inguinal lymph node after two weeks of cyclic tibial compression.** Comparison of the number of CD3+, CD3+CD4+Tbet+, CD3+CD4+Rorγt+, and CD3+CD4+Rorγt+Tbet+ T cells in the inguinal lymph node of loaded and contralateral limbs of

16-week old female saline and fingolimod treated C57Bl/6 mice after two weeks of loading. *p<.05

Supplementary Figure S4

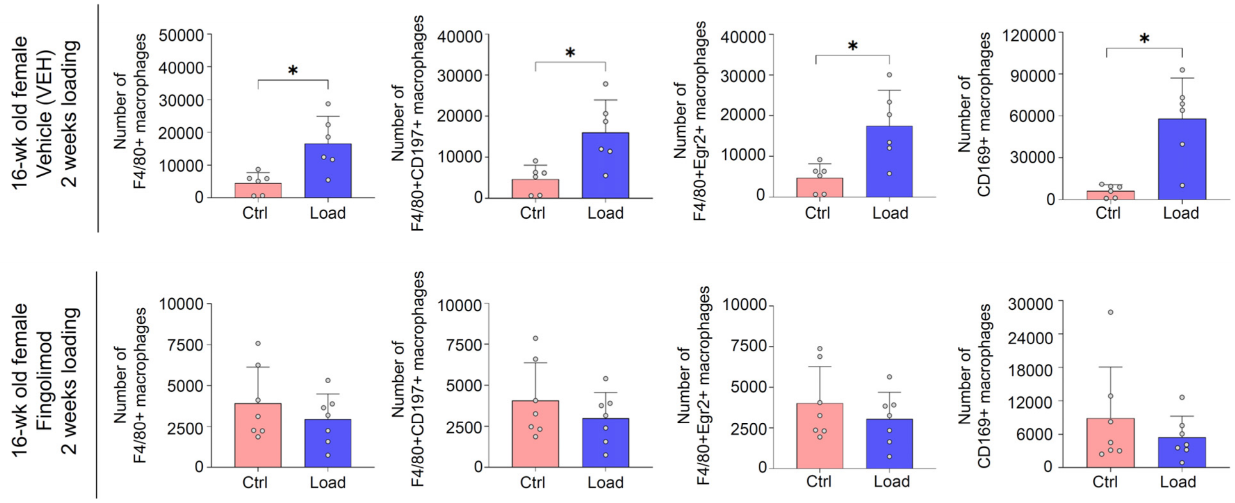


**Supplementary Figure S4, Related to Figure 4. Fingolimod treatment did not change the number of macrophages in the inguinal lymph node after two weeks of cyclic tibial compression.** Comparison of the number of F4/80+, F480+CD197+, F4/80+Egr2+, and CD169+ macrophages in the inguinal lymph node of loaded and contralateral limbs of 16-week old female saline and fingolimod treated C57Bl/6 mice after two weeks of loading. *p<.05

Supplementary Figure S5

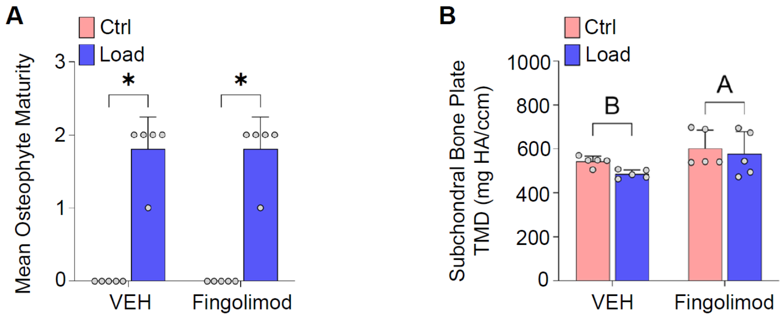


**Supplementary Figure S5, Related to Figure 4. Fingolimod treatment did not attenuate osteophyte maturity after two weeks of cyclic tibial compression. A)** Comparison of mean osteophyte maturity in the medial tibial plateau in 16-week old female saline and fingolimod treated C57Bl/6 mice after two weeks of loading. *p<.05 **B)** microCT analysis comparison of subchondral bone plate TMD after two weeks of loading and saline or fingolimod treatment. A>B>C

Supplementary Figure S6

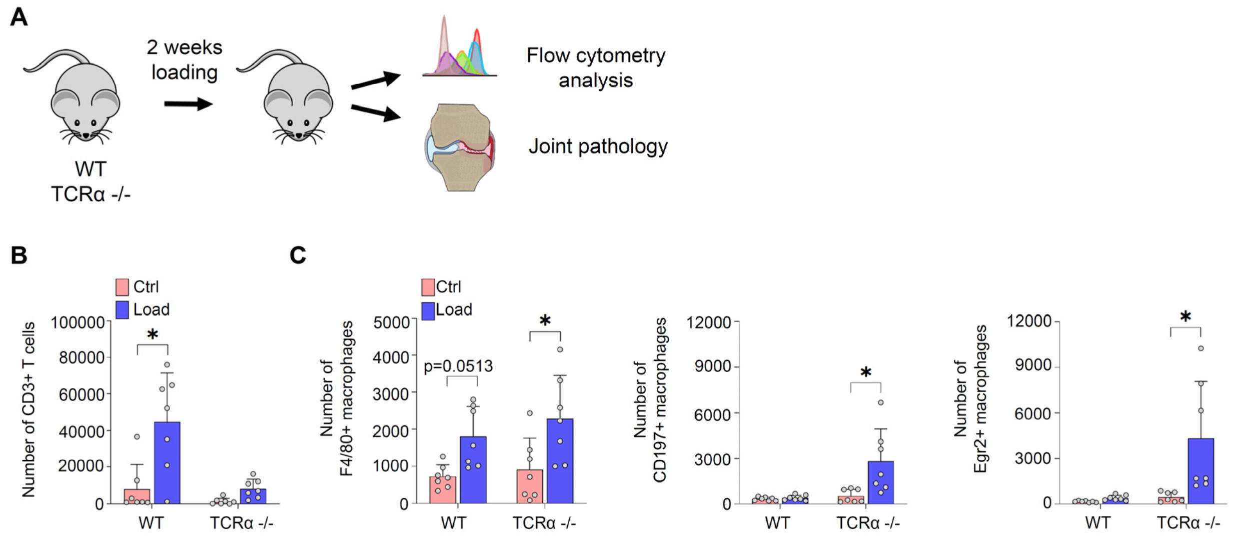


**Supplementary Figure S6, Related to Figure 5. The number of macrophages increased in the inguinal lymph node in the absence of αβ T cells after two weeks of cyclic tibial compression.** **A)** Schematic of experimental design. 16-week old female WT and TCRα -/- mice underwent two weeks of cyclic tibial compression. The inguinal lymph node was assessed through flow cytometry and joints were assessed through histology and microCT. **B)** Comparison of the number of CD3+ T cells in the inguinal lymph node of loaded and contralateral limbs of 16-week old female WT and TCRα -/- mice after two weeks of loading. **C)** Comparison of the number of F4/80+ (pan marker), CD197+ (M1), and Egr2+ (M2) macrophages in the inguinal lymph node of loaded and contralateral limbs of 16-week old female WT and TCRα -/- mice after two weeks of loading. *p<.05

Supplementary Figure S7

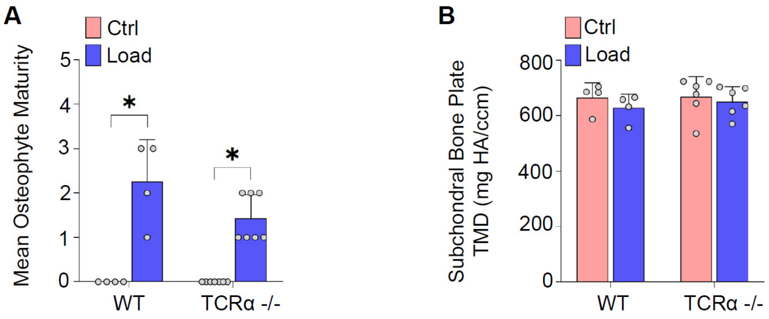


**Supplementary Figure S7, Related to Figure 5. The absence of αβ T cells did not attenuate osteophyte maturity. A)** Comparison of mean osteophyte maturity in the medial tibial plateau in 16-week old female C57Bl/6 and TCRα -/- mice after two weeks of loading. *p<.05 **B)** Comparison of subchondral plate tissue mineral density in 16-week old female C57Bl/6 and TCRα -/- mice after two weeks of loading.

**Supplementary Table 1. Surface and Intracellular Markers for Flow Cytometry.** Manufacturer and clone information for surface and intracellular markers used for flow cytometry.

| Marker | Clone | Manufacturer |
|---|---|---|
| APC-anti-CD3 | 17A2 | eBioscience |
| FITC-anti-CD4 | RM4-5 | eBioscience |
| PE-Cy7-anti-CD8a | 53-6.7 | eBioscience |
| FITC-anti-γδTCR | GL3 | eBioscience |
| PE-anti-F4/80 | BM8 | eBioscience |
| PE-Cy7-anti-CD197 | 4B12 | eBioscience |
| APC-anti-Egr2 | erongr2 | eBioscience |
| PE-Cy7-anti-CD19 | 1D3 | eBioscience |
| Alexa Fluor 488-anti-FoxP3 | FJK-16s | eBioscience |
| PE-Cy7-anti-Rorγt | B2D | eBioscience |
| APC-anti-γδTCR | GL3 | eBioscience |
| PE-anti-CD169 | SER-4 | eBioscience |
| Alexa Fluor 488-anti-Egr2 | erongr2 | eBioscience |
| BUV395-anti-CD3 | 17A2 | BD Biosciences |
| BV510-anti-CD4 | RM4-5 | BD Biosciences |
| BUV737-anti-CD8a | 53-6.7 | BD Biosciences |
| BV650-anti-Tbet | O4-46 | BD Biosciences |
| BV421-anti-GATA3 | L50-823 | BD Biosciences |
| BUV395-anti-F4/80 | T45-2342 | BD Biosciences |
| BV605-anti-CD197 | 4B12 | BD Biosciences |
